# Human Cytomegalovirus Genomes Sequenced Directly from Clinical Material: Variation, Multiple-Strain Infection, Recombination and Mutation

**DOI:** 10.1101/505735

**Authors:** Nicolás M. Suárez, Gavin S. Wilkie, Elias Hage, Salvatore Camiolo, Marylouisa Holton, Joseph Hughes, Maha Maabar, Vattipally B. Sreenu, Akshay Dhingra, Ursula A. Gompels, Gavin W. G. Wilkinson, Fausto Baldanti, Milena Furione, Daniele Lilleri, Alessia Arossa, Tina Ganzenmueller, Giuseppe Gerna, Petr Hubáček, Thomas F. Schulz, Dana Wolf, Maurizio Zavattoni, Andrew J. Davison

## Abstract

The genomic characteristics of human cytomegalovirus (HCMV) strains sequenced directly from clinical pathology samples were investigated, focusing on variation, multiple-strain infection, recombination and natural mutation. A total of 207 datasets generated in this and previous studies using target enrichment and high-throughput sequencing were analysed, in the process facilitating the determination of genome sequences for 91 strains. Key findings were that (i) it is important to monitor the quality of sequencing libraries in investigating diversity, (ii) many recombinant strains have been transmitted during HCMV evolution, and some have apparently survived for thousands of years without further recombination, (iii) mutants with non-functional genes (pseudogenes) have been circulating and recombining for long periods and can cause congenital infection and resulting clinical sequelae, and (iv) intrahost diversity in single-strain infections is much less than that in multiple-strain infections. Future population-based studies are likely to continue illuminating the evolution, epidemiology and pathogenesis of HCMV.

## BACKGROUND

Human cytomegalovirus (HCMV) poses a risk particularly to people with immature or compromised immune systems, and can have serious outcomes in congenitally infected children, transplant recipients and people affected by HIV/AIDS. Prior to the advent of high-throughput technologies, studies of HCMV genomes in natural infections were limited to Sanger sequencing of PCR amplicons, often focusing on a small number of polymorphic (hypervariable) genes [1]. This left out most of the genome and also restricted the characterisation of multiple-strain infections, which may have more serious outcomes.

The first complete HCMV genome sequence to be determined was that of the high-passage strain AD169 [2], from a plasmid library. Over a decade later, additional genomes were sequenced from bacterial artificial chromosomes [3–5], virion DNA [6] or overlapping PCR amplicons [7, 8]. These sequences were determined using Sanger technology, and were complemented subsequently by many others, increasingly using high-throughput methods [7, 9–13]. With only three exceptions [7, 11], all were derived from laboratory strains isolated in cell culture. Mounting evidence of the existence of multiple-strain infections and the propensity of HCMV to mutate during cell culture [6, 7, 8, 14, 15] added impetus to sequencing genomes directly from clinical material in order to define natural populations. One strategy for this involves sequencing overlapping PCR amplicons [7, 16]. Another utilises an oligonucleotide bait library representing known HCMV diversity to select target sequences from random DNA fragments. This target enrichment technology originated in commercial kits for cellular exome sequencing, and was subsequently applied to various pathogens [17, 18], including HCMV [19–21]. We have applied it since 2012 and have systematically released via GenBank many genome sequences that have proved pivotal in other studies [11, 12, 19–21].

The HCMV genome exhibits several evolutionary phenomena, including hypervariation, multiple-strain infection, recombination and gene loss by mutation, all of which were discovered prior to high-throughput sequencing and have since been illuminated by this technology (early references are [22–26]). In this article, we explore these and other key genomic features of HCMV, with an emphasis on the strains present in clinical material.

## METHODS

### Samples

For convenience, samples were analysed as three collections detailed in **Supplementary Tables 1-3** and summarised in Table 1, which also contains information on ethical permissions. Collection 1 originated from congenital infections from Pavia, Jerusalem and Prague. Collection 2 originated from Hannover and Pavia, and most came from transplant recipients. Collection 3 represents samples obtained by others in previous studies from people with various conditions and sequenced using target enrichment, but with a different oligonucleotide bait library. The features of the samples are in **Supplementary Tables 1-3** rows 3-6, and the clinical outcomes of congenital infection are in **Supplementary Table 1** row 205.

**Table 1.**
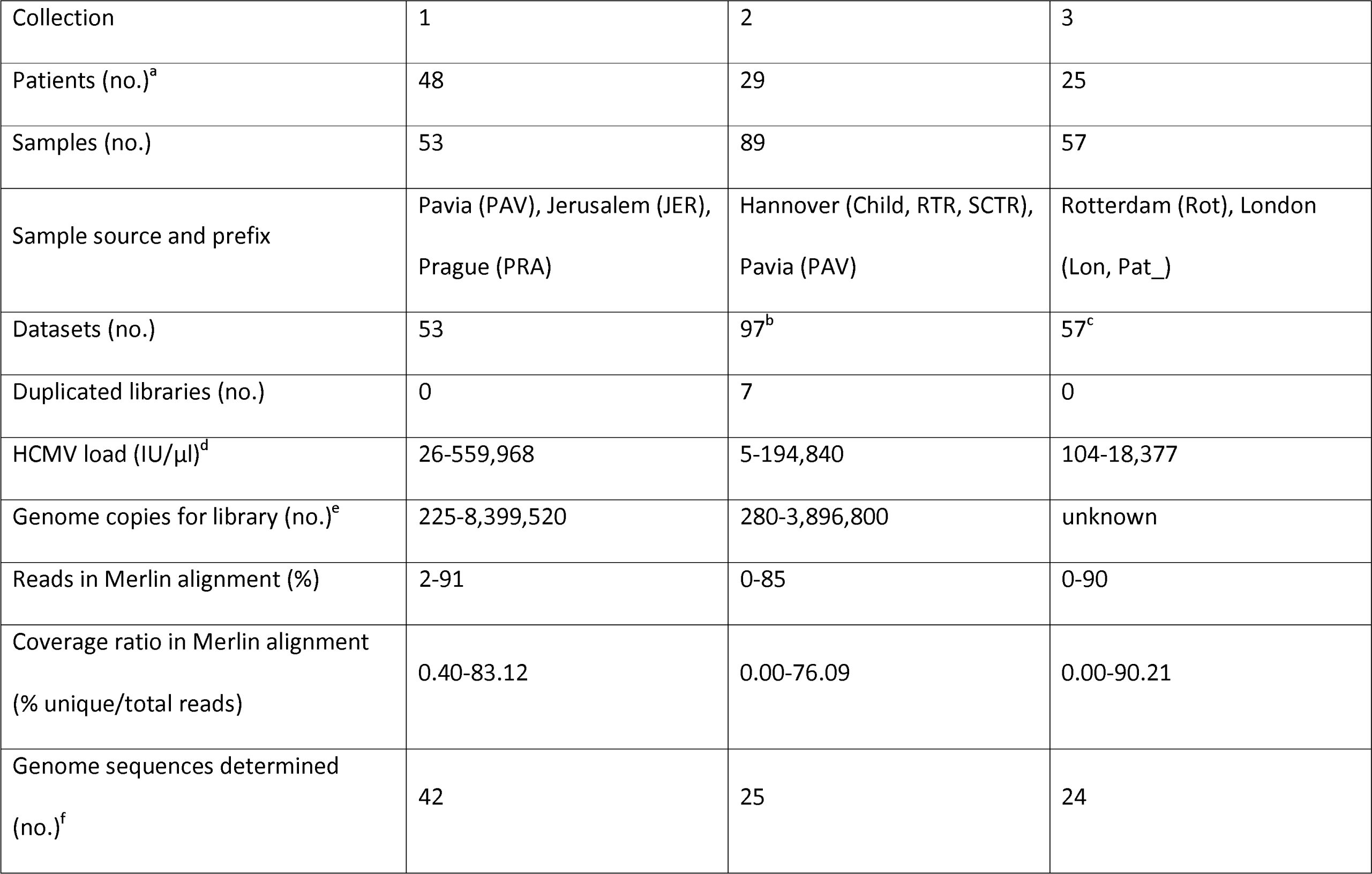
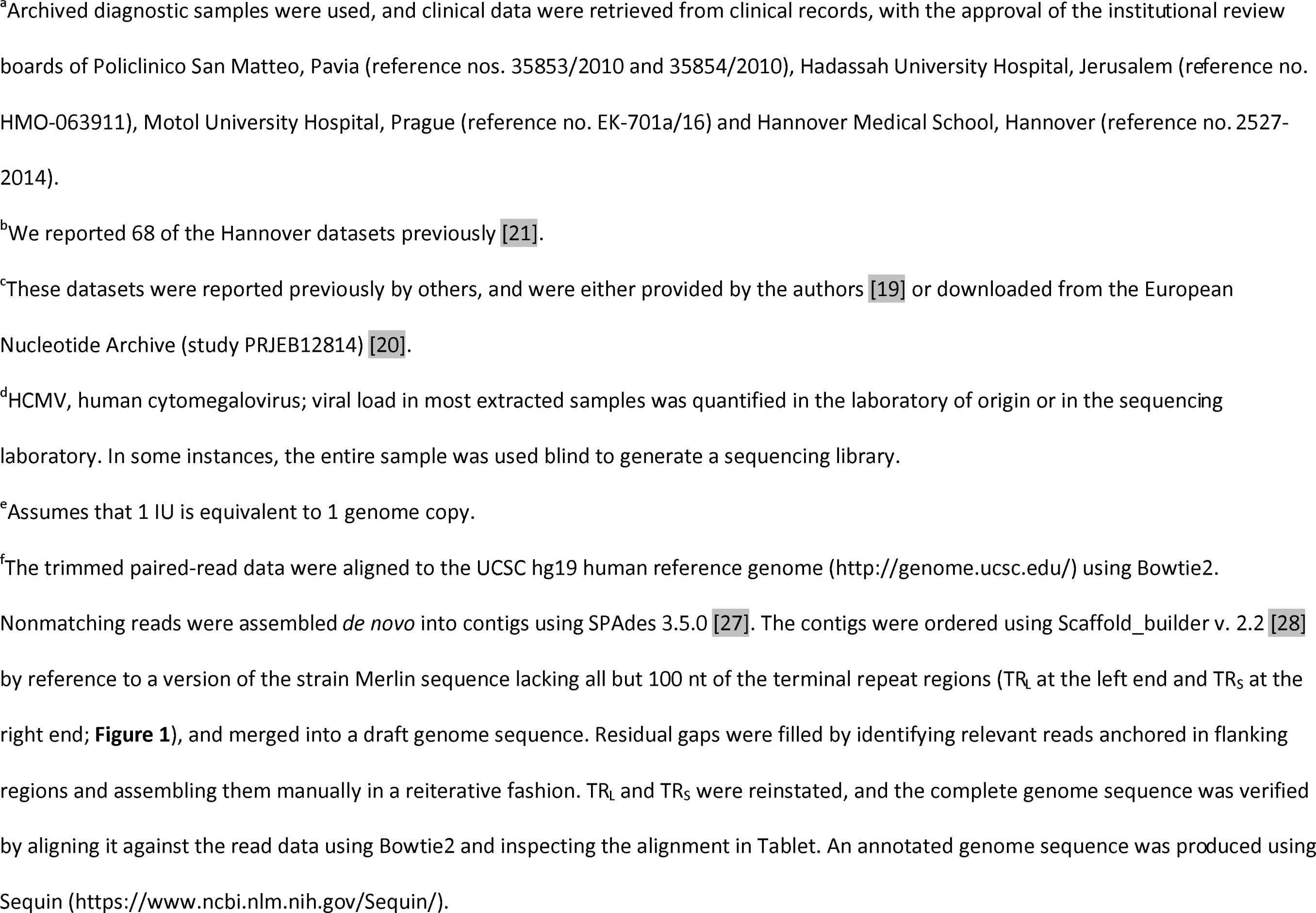
Selected information on the sample collections. Full details are provided in **Supplementary Tables 1-3**.

| Collection | 1 | 2 | 3 |
| --- | --- | --- | --- |
| Patients (no.)<sup>a</sup> | 48 | 29 | 25 |
| Samples (no.) | 53 | 89 | 57 |
| Sample source and prefix | Pavia (PAV), Jerusalem (JER),<br>Prague (PRA) | Hannover (Child, RTR, SCTR),<br>Pavia (PAV) | Rotterdam (Rot), London<br>(Lon, Pat_) |
| Datasets (no.) | 53 | 97<sup>b</sup> | 57<sup>c</sup> |
| Duplicated libraries (no.) | 0 | 7 | 0 |
| HCMV load (IU/μl)<sup>d</sup> | 26-559,968 | 5-194,840 | 104-18,377 |
| Genome copies for library (no.)<sup>e</sup> | 225-8,399,520 | 280-3,896,800 | unknown |
| Reads in Merlin alignment (%) | 2-91 | 0-85 | 0-90 |
| Coverage ratio in Merlin alignment<br>(% unique/total reads) | 0.40-83.12 | 0.00-76.09 | 0.00-90.21 |
| Genome sequences determined<br>(no.)<sup>f</sup> | 42 | 25 | 24 |

<sup>a</sup>Archived diagnostic samples were used, and clinical data were retrieved from clinical records, with the approval of the institutional review boards of Policlinico San Matteo, Pavia (reference nos. 35853/2010 and 35854/2010), Hadassah University Hospital, Jerusalem (reference no. HMO-063911), Motol University Hospital, Prague (reference no. EK-701a/16) and Hannover Medical School, Hannover (reference no. 2527-2014).
<sup>b</sup>We reported 68 of the Hannover datasets previously [21].
<sup>c</sup>These datasets were reported previously by others, and were either provided by the authors [19] or downloaded from the European Nucleotide Archive (study PRJEB12814) [20].
<sup>d</sup>HCMV, human cytomegalovirus; viral load in most extracted samples was quantified in the laboratory of origin or in the sequencing laboratory. In some instances, the entire sample was used blind to generate a sequencing library.
<sup>e</sup>Assumes that 1 IU is equivalent to 1 genome copy.
<sup>f</sup>The trimmed paired-read data were aligned to the UCSC hg19 human reference genome (http://genome.ucsc.edu/) using Bowtie2. Nonmatching reads were assembled *de novo* into contigs using SPAdes 3.5.0 [27]. The contigs were ordered using Scaffold\_builder v. 2.2 [28] by reference to a version of the strain Merlin sequence lacking all but 100 nt of the terminal repeat regions (TR<sub>L</sub> at the left end and TR<sub>S</sub> at the right end; **Figure 1**), and merged into a draft genome sequence. Residual gaps were filled by identifying relevant reads anchored in flanking regions and assembling them manually in a reiterative fashion. TR<sub>L</sub> and TR<sub>S</sub> were reinstated, and the complete genome sequence was verified by aligning it against the read data using Bowtie2 and inspecting the alignment in Tablet. An annotated genome sequence was produced using Sequin (https://www.ncbi.nlm.nih.gov/Sequin/).

### DNA sequencing

Target enrichment and sequencing library preparation were performed using the SureSelect^XT^ v. 1.7 system for Illumina paired-end libraries with biotinylated cRNA probe baits (Agilent, Stockport, UK) [21]. Bait libraries representing known HCMV diversity were designed in February 2012 and April 2014 from 31 and 64 complete genome sequences, respectively. Access to the latter library is available from the corresponding author. Data on viral loads and library construction are in **Supplementary Tables 1-3** rows 9-12. Datasets of 300 or 150 nt paired-end reads were generated using a MiSeq (Illumina, San Diego, CA, USA). Their names are in **Supplementary Tables 1-3** row 7. They were prepared for analysis using Trim Galore v 0.4.0 (http://www.bioinformatics.babraham.ac.uk/projects/trim_galore/; length=21, quality=10 and stringency=3). The numbers of trimmed reads are in **Supplementary Tables 1-3** row 15.

### Library diversity

The number of reads in a dataset that were derived from unique HCMV fragments was estimated by using Bowtie2 v. 2.2.6 [29] to align the reads against the strain Merlin sequence (GenBank accession AY446894.2), and also, where it could be determined, the consensus genome sequence derived from the dataset. The relevant data are in **Supplementary Tables 1-3** rows 17-19 and 23-26. Reads containing insertions or deletions, duplicate read pairs sharing both end coordinates and duplicate unpaired reads sharing one end coordinate were removed, thereby producing an alignment file for unique reads derived from unique HCMV fragments (https://centre-for-virus-research.github.io/VATK/AssemblyPostProcessing). This file was viewed using Tablet v. 1.14.11.7 [30]. The coverage depth values for total and unique fragment reads are in **Supplementary Tables 1-3 rows 20-21 and 27-28**.

### Strain enumeration

The numbers of strains represented in each dataset were estimated by two strategies: genotype read-matching and motif read-matching (https://centre-for-virus-research.github.io/VATK/HCMV_pipeline). Both strategies utilised datasets concatenated from the paired-end datasets. The genotype designations used were either based on reported phylogenies [6, 12, 25, 31, 32], amended or extended as appropriate, or were constructed afresh using Clustal Omega v. 1.2.4 [33] and MEGA v. 6.0.6 [34] with data for the genomes listed in **Supplementary Table 4** and individual genes for which additional sequences were available in GenBank. Alignments are in **Supplementary Figure 1**.

For genotype read-matching, Bowtie2 was used to align the reads to sequences representing the genotypes of two hypervariable genes, UL146 and RL13 [6, 12, 35]. The sequences representinng the entire coding region of UL146 and the central coding region of RL13 are in **Supplementary Tables 1-3** rows 34-58. In contrast to the UL146 genotypes, the RL13 genotypes cross-matched to some extent within four groups (G1, G2, G3; G4A, G4B; G6, G10; and G7, G8). In these instances, the genotype within the group with most matching reads was scored. The numbers of reads aligned to each genotype are in **Supplementary Tables 1-3** rows 34-58. A genotype was scored if the number of reads was >10 and represented >2% of the total number detected for all genotypes of that gene. For 14 samples in collection 1 that had been sequenced prior to the availability of ultrapure (TruGrade) oligonucleotides, these values were set at >25 and >5%, respectively. The number of strains in a sample was scored as the greater of the numbers of genotypes detected for the two target genes, and is in **Supplementary Tables 1-3** row 13.

For motif read-matching, conserved genotype-specific motifs (20-31 nt) were identified by visual inspection of the alignments for 12 hypervariable genes (**Supplementary Figure 1**) [6, 12, 19, 33]. Additional motifs were included in order to identify the more common intergenotypic recombinants and pseudogenic mutants. The motif sequences and number of reads containing perfect matches to a sequence or its reverse complement are in **Supplementary Tables 1-3** rows 60-178. Genotypes were scored as described above. The number of strains in a sample was scored as the maximum number of genotypes detected for at least two genes, and is in **Supplementary Tables 1-3** row 14.

### Data deposition

The original read datasets were purged of human data and then deposited in the European Nucleotide Archive (ENA; project no. PRJEB29585), and the consensus genome sequences were deposited in GenBank. The accession numbers are in **Supplementary Tables 1-3** rows 8 and 29, respectively. Updated genome sequence determinations in collection 3 were deposited by the original submitters in GenBank [19] or by us as third-party annotations in ENA (project no. PRJEB29374) [20]. Sequence features are in **Supplementary Tables 1-3** rows 30-32. Data on pseudogenic mutants (obtained by motif read-matching) in collection 1 are in **Supplementary Table 1** rows 180-203.

### Intrahost variant analysis

Minor variant nucleotides were examined in datasets for which a consensus genome sequence had been determined. Original datasets were prepared for analysis using Trim Galore (length=100, quality=30 and stringency=1), and trimmed reads were mapped using Bowtie2. Alignment files in SAM format were converted into BAM format, sorted using SAMtools v. 1.3 [36], and analysed using LoFreq v. 2.1.2 [37] and V-Phaser 2 [38] under default parameters.

## RESULTS

### Operational limitations

A total of 207 datasets from 199 samples and 102 individuals were analysed (Table 1 and **Supplementary Tables 1-3**). Library quality was represented in the percentage of HCMV reads and the coverage depth by unique fragment reads. These values were related to sample type, being higher for urine than blood presumably because of a higher proportion of viral to host DNA. They also depended on the number of viral genome copies used to make the library, with >1000 copies generally being needed to determine a complete genome sequence. However, despite high library diversity, it was not possible to assemble complete genome sequences from most datasets in collection 3 because of gaps in RL12 and G+C-rich regions. This may have been because of limitations in the bait library and sequencing approach used. The use of excessive PCR cycles with some samples in collections 1 and 2 led to high coverage depth by total fragment reads but low coverage depth by unique fragment reads, and thus to highly clonal libraries (e.g. PAV2 in collection 1). Genotypes present at subthreshold levels may represent multiple-strain infections or cross-contamination during the complex sample processing pathway (e.g. PRA4 reads in PRA6 dataset 78943 in collection 1).

### Genome sequences

A total of 91 complete or almost complete HCMV genome sequences were determined (Table 1). We reported five previously [21], and 16 are improvements on published sequences [19]. Most originated from single-strain infections or multiple-strain infections in which one strain was predominant, and some originated from different strains that predominated in a patient at different times. Defining a strain on the basis of a viral genome present in an individual, these 91 sequences, plus an additional 49 deposited by our group and 104 by other groups, brought the number of strains sequenced to 244 (**Supplementary Table 4**). Of these, 91 were sequenced directly from clinical material, and all but one were determined in this and our previous report [21]. The average size of the HCMV genome, based on the 78 complete sequences in this set, is 235,465 bp (range 234,316-237,120 bp).

### Multiple-strain infections

Genotypic differences in hypervariable genes (Figure 1 and **Supplementary Figure 1**) were exploited to detect multiple-strain infections by genotype read-matching and motif read-matching, the latter method proving to be the more sensitive. Single strains were common in congenitally infected patients (n=43/50 in collections 1 and 2), but significantly less so in transplant recipients (n=11/25 in collections 2 and 3; chi-square test, χ^2^=14.583, *p*<0.05).

**Figure 1.**
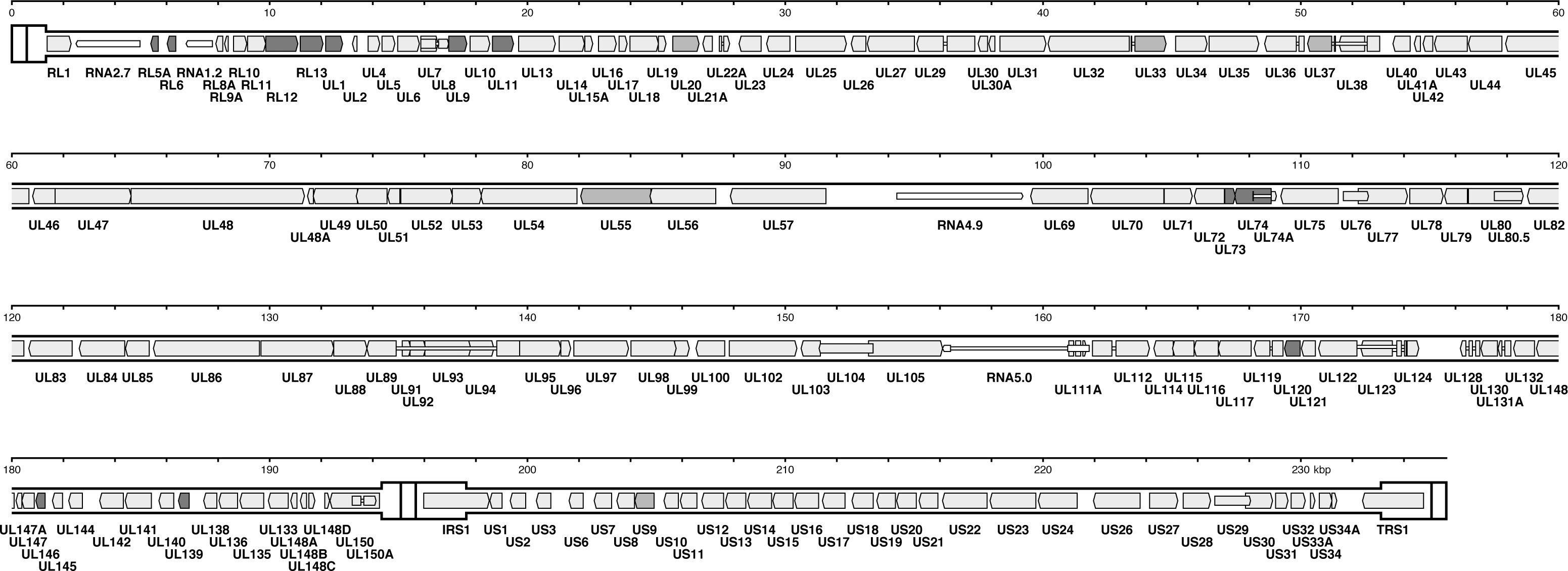
Locations in the human cytomegalovirus strain Merlin genome of genes used for genotyping. The genome consists of two unique regions, U_L_ (1325-194343 bp) and U_S_ (197627-233108 bp), the former flanked by inverted repeats TR_L_ (1-1324 bp) and IR_L_ (194344-195667 bp), and the latter flanked by inverted repeats IR_S_ (195090-197626 bp) and TR_S_ (233109-235646 bp). Protein-coding regions are indicated by colour-shaded arrows, and noncoding RNAs as narrower, white-shaded arrows, with gene nomenclature below. Introns are shown as narrow white bars. The 12 genes (RL5A, RL6, RL12, RL13, UL1, UL9, UL11, UL73, UL74, UL120, UL146 and UL139) used for motif read-matching are coloured red < dark *grey in greyscale version* >. Two of these genes (RL13 and UL146) were also used for genotype read-matching. The additional five genes (UL20, UL33, UL37, UL55 and US9) used to genotype genome sequences by alignment are coloured orange <mid *grey in greyscale version*>. All other genes are coloured pink <light *grey in greyscale version*>.

### Recombination

The 244 genome sequences were genotyped in the 12 hypervariable genes used for motif read-matching and then in five additional genes (Figure 1 and **Supplementary Table 4**).

Hypervariation in UL55, which encodes glycoprotein B (gB), is located in two regions (UL55N, near the N terminus, and UL55X, encompassing the proteolytic cleavage site) [23, 39]. Five genotypes (G1-G5) have been assigned to each region [23, 39-41], which are separated by 927 bp that are 80% identical in all strains. All genomes had one of the recognised UL55X genotypes (**Supplementary Table 5**). As reported previously [39], UL55N G2 and G3 could not be distinguished reliably from each other, and two additional genotypes (G6-G7) were detected that may have arisen from ancient recombination events within UL55N (**Supplementary Tables 4 and 5**, and **Supplementary Figure 1**). There was evidence for recombination in the region between UL55N and UL55X in only eight genomes. This low proportion of recombination (3.3%) contrasts with the higher levels proposed previously from PCR-based studies [39, 42].

UL73 and UL74, which encode glycoproteins N and O (gN and gO), respectively, are adjacent hypervariable genes that exist as eight genotypes each [25, 32, 43]. There was evidence for recombination between them in only seven genomes (2.9%), in accordance with the low levels (2.2%) first detected in PCR-based studies [25, 32, 44]. In the region containing adjacent hypervariable genes RL12, RL13 and UL1, recombinants were also rare (1.2%) within RL12 and absent from RL13 and UL1. In contrast, hypervariable genes UL146 and UL139, which encode a CXC chemokine and a membrane glycoprotein, respectively, are separated by a well-conserved region of over 5 kbp. The number (66) of the 126 possible genotype combinations represented in the 244 genomes is too large to allow any underlying genotypic linkage to be discerned, consistent with previous conclusions from PCR-based studies [31]. No recombinants were noted within UL146.

In principle, strains in multiple-strain infections have the opportunity to recombine. In our previous analysis of RTR1 in collection 2, we noted that one strain (RTR1A) predominated at earlier times and another (RTR1B) at later times [21]. From the low frequency of variants across a large part of the genome, we concluded that the second strain had arisen either by recombination involving the first strain or by reinfection with, or reactivation of, a second strain fortuitously similar to the first. In the present study, recombination was strongly supported by a comparison of the two genome sequences, which showed that approximately two-thirds of the genome is almost identical (differing by three substitutions in noncoding regions), whereas the remaining third is highly dissimilar.

To investigate whether strains have been transmitted without recombination occurring, identical genotypic constellations were identified among the 244 genomes (**Supplementary Table 6**). This revealed the existence of 12 haplotype groups within which multiple strains exhibit no signs of having recombined since diverging from their last common ancestor; these are termed nonrecombinant strains below. As an incidental outcome, the two strains in group 1 (PRA8 and CZ/3/2012), which were characterised in different studies, were confirmed as having originated from the same patient, reducing the set of sequenced strains to 243. The results from the other 11 groups suggest that nonrecombinant strains have been circulating, some for periods sufficient to allow the accumulation of >100 substitutions. Application of an evolutionary rate estimated for herpesviruses (3.5 = 10^−8^ substitutions/nt/year) [45] implies that these periods may have extended to many thousands of years. The distribution of substitutions across the genome in highly divergent groups 9 and 10 was examined in further detail. Group 9 (three strains) exhibited 135 differences, with the 50 that would affect protein coding distributed among 38 genes, and group 10 (two strains) exhibited 138 differences, with the 38 that would affect protein coding distributed among 27 genes. No obvious bias was observed towards greater diversity in any particular gene or group of genes, including those in the hypervariable category. As suggested by the lack of diversity within genotypes in comparison with the marked diversity among them (**Supplementary Figure 1**), these findings fit with the view that intense diversification of the hypervariable genes occurred early in human or pre-human history [25, 31] and has long since ceased.

### Pseudogenes

Mutations have been catalogued in some HCMV strains that cause premature translational termination [7, 11, 12, 26]. They take the form of substitutions introducing in-frame stop codons or ablating splice sites, or insertions, deletions or inversions causing frameshifting or loss of entire protein-coding regions. However, evidence for these viral pseudogenes was derived mostly from strains isolated in cell culture, and it was unclear to what extent they are present in natural populations. For example, in a study reporting that 75% of strains are mutated [12], 157 mutations were identified in 101 strains, with all but one passaged in cell culture and 35 confirmed by PCR of the clinical material. Nonetheless, the distribution of mutations among the 91 strains sequenced in the present study from clinical material is similar to that among strains isolated in cell culture (Table 2 and **Supplementary Table 4**), thus validating the earlier conclusions.

**Table 2.** Mutated genes in order of decreasing frequency.

| Gene | Strains mutated (no.)<sup>a</sup> |  |  | Strains mutated (%)<sup>a</sup> |  |  |
| --- | --- | --- | --- | --- | --- | --- |
|  | Passaged<sup>b</sup> | Clinical<sup>c</sup> | All<sup>d</sup> | Passaged<sup>b</sup> | Clinical<sup>c</sup> | All<sup>d</sup> |
| UL9 | 50 | 31 | 81 | 32.89 | 34.07 | 33.33 |
| RL5A | 31 | 27 | 58 | 20.39 | 29.67 | 23.87 |
| UL1 | 20 | 18 | 38 | 13.16 | 19.78 | 15.64 |
| RL6 | 23 | 14 | 37 | 15.13 | 15.38 | 15.23 |
| US9 | 26 | 11 | 37 | 17.11 | 12.09 | 15.23 |
| UL111A | 16 | 7 | 23 | 10.53 | 7.69 | 9.47 |
| UL150 | 11 | 3 | 14 | 7.24 | 3.30 | 5.76 |
| US7 | 7 | 7 | 14 | 4.61 | 7.69 | 5.76 |
| UL40 | 8 | 2 | 10 | 5.26 | 2.20 | 4.12 |
| UL30 | 2 | 3 | 5 | 1.32 | 3.30 | 2.06 |
| UL142 | 2 | 3 | 5 | 1.32 | 3.30 | 2.06 |
| RL12 | 3 | 1 | 4 | 1.97 | 1.10 | 1.65 |
| RL1 | 1 | 2 | 3 | 0.66 | 2.20 | 1.23 |
| UL136 | 3 | 0 | 3 | 1.97 | 0.00 | 1.23 |
| US13 | 3 | 0 | 3 | 1.97 | 0.00 | 1.23 |
| UL133 | 2 | 0 | 2 | 1.32 | 0.00 | 0.82 |
| US6 | 1 | 1 | 2 | 0.66 | 1.10 | 0.82 |
| US8 | 0 | 2 | 2 | 0.00 | 2.20 | 0.82 |
| US27 | 2 | 0 | 2 | 1.32 | 0.00 | 0.82 |
| UL11 | 1 | 0 | 1 | 0.66 | 0.00 | 0.41 |
| UL13 | 0 | 1 | 1 | 0.00 | 1.10 | 0.41 |
| UL14 | 0 | 1 | 1 | 0.00 | 1.10 | 0.41 |
| UL15A | 0 | 1 | 1 | 0.00 | 1.10 | 0.41 |
| UL20 | 1 | 0 | 1 | 0.66 | 0.00 | 0.41 |
| UL43 | 0 | 1 | 1 | 0.00 | 1.10 | 0.41 |
| UL99 | 1 | 0 | 1 | 0.66 | 0.00 | 0.41 |
| UL148 | 1 | 0 | 1 | 0.66 | 0.00 | 0.41 |
| UL147 | 1 | 0 | 1 | 0.66 | 0.00 | 0.41 |
| UL145 | 0 | 1 | 1 | 0.00 | 1.10 | 0.41 |
| UL150A | 1 | 0 | 1 | 0.66 | 0.00 | 0.41 |
| IRS1 | 1 | 0 | 1 | 0.66 | 0.00 | 0.41 |
| US1 | 1 | 0 | 1 | 0.66 | 0.00 | 0.41 |
| US12 | 1 | 0 | 1 | 0.66 | 0.00 | 0.41 |
| US19 | 0 | 1 | 1 | 0.00 | 1.10 | 0.41 |

<sup>a</sup>Omitting mutations that occurred in RL13, UL128, UL130 and UL131A probably during passage, or that were engineered during bacterial artificial chromosome construction.
<sup>b</sup>Strains sequenced from strains passaged in cell culture, not taking into account the minority of mutations confirmed from the clinical samples (n=152, excludes CZ/3/2012, which is the same strain as PRA8).
<sup>c</sup>Strains sequenced directly from clinical material (n=91).
<sup>d</sup>Strains sequenced directly from clinical material or passaged virus (n=243).

Among the strains sequenced from clinical material, 77% are mutated in at least one gene (compared with 79% among all sequenced strains), and one is mutated in as many as six genes (Pat_D in collection 3). The most frequently mutated genes are UL9, RL5A, UL1 and RL6 (members of the RL11 family), US7 and US9 (members of the US6 gene family), and UL111A (encoding viral interleukin-10) (**Supplementary Figure 1**). The likelihood that many of these mutants are ancient is supported by the finding that all were detected at levels very close to 100% in collection 1, and by previous observations identifying the same mutation in different strains [7, 12]. In addition, there was evidence from the PAV6 datasets for maternal transmission of a US7 mutant (**Supplementary Table 1**), and from PCR data (not shown) for maternal transmission of a UL111A mutant to PAV16. Moreover, nine of the groups of nonrecombinant strains contained mutants, and some of the mutations were common to group members (**Supplementary Table 6**) and even to additional strains among the 243, indicating that they had been transferred by recombination. These observations again indicate the longevity of many of the pseudogenes and their propagation by recombination. Focusing on the most common mutations, strains in which UL9, RL5A, UL1, US9, US7 and UL111A were affected (singly or in combination) were, like strains that are not mutated in any gene, transmitted in congenital infection and, in some cases, linked to defects in neurological development (**Supplementary Table 1**).

### Intrahost diversity

In addition to mutations indicative of pseudogenes, HCMV populations in individuals may include minor variant nucleotides (single nucleotide polymorphisms). LoFreq and V-Phaser analyses showed that single-strain infections contained markedly fewer variants (median values of 60 and 140, respectively) than multiple-strain infections (median values of 2444 and 2955, respectively; Figure 2). The differences between the values for single- and multiple-strain infections were significant (Kruskal-Wallis rank-sum test; LoFreq, χ^2^=67.918, *p*<2.2e^−16^; V-Phaser, χ^2^=63.536, *p*=1.6e^−15^). Seven outliers in single-strain infections were common to both analyses (in order of decreasing number of variants, RTR6B, CMV-37, RTR2, CMV-35, CMV-38, ERR1279054 and PAV6), one was reported by LoFreq only (PAV21), and four were reported by V-Phaser only (CMV-19, CMV-31, PRA6A and SCRT12).

**Figure 2.**
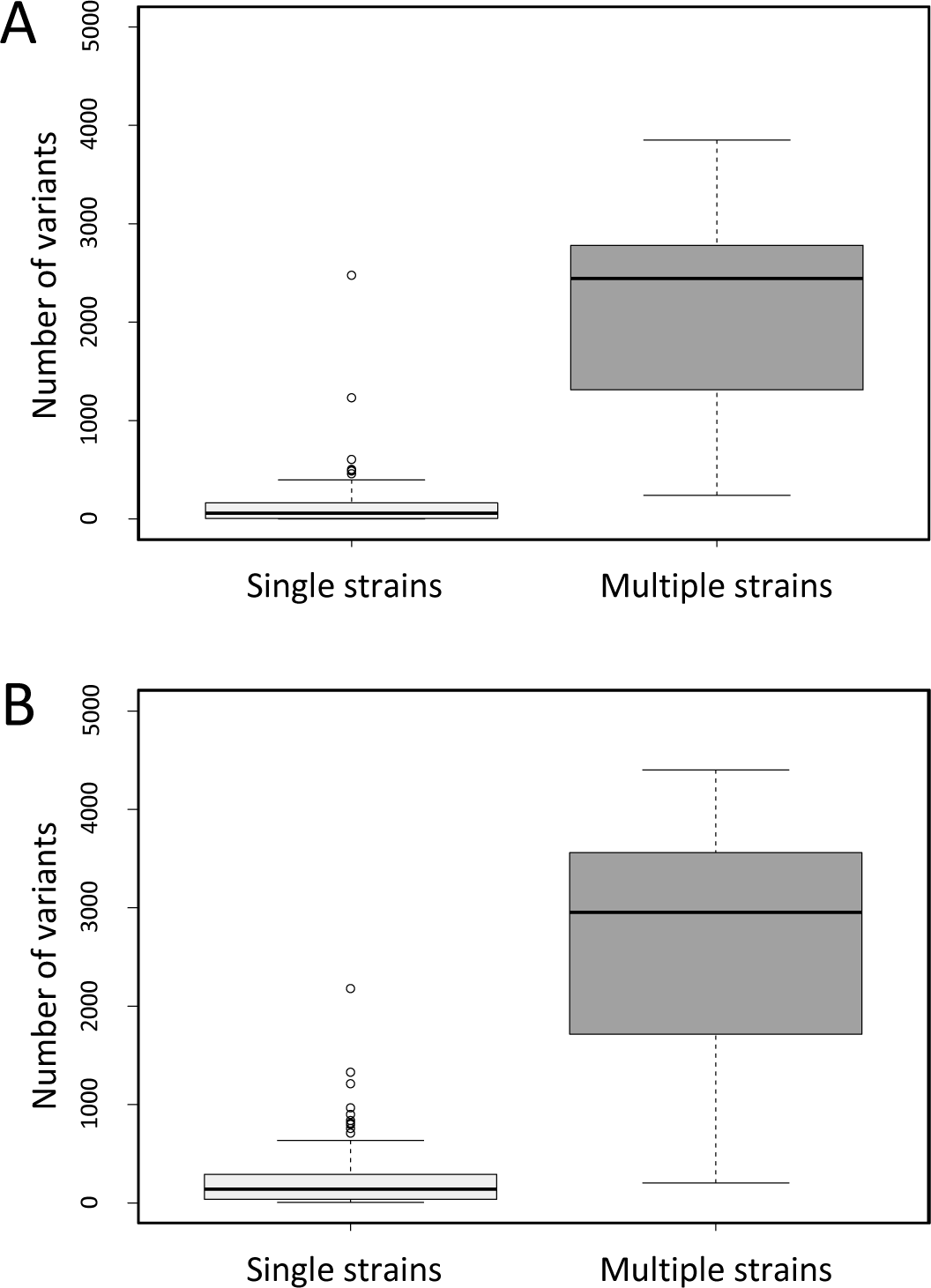
Box-and-whisker graphs showing the number of variants detected in single-strain and multiple-strain infections using (A) LoFreq and (B) V-Phaser. The graphs were created using ggplot2 (https://ggplot2.tidyverse.org). Single-strain (n=134 and 131, respectively) and multiple-strain datasets (n=29 and 29, respectively) for which consensus genome sequences were derived were identified by motif read-matching. The total number of variants (2-50% of the total) in each dataset was enumerated; length polymorphisms were not considered. Each box (light grey for single strains and dark grey for multiple strains) encompasses the first to third quartiles (Q1 to Q3) and shows the median as a thick line. For each box, the horizontal line at the end of the upper dashed whisker marks the upper extreme (defined as the smaller of Q3+1.5(Q3-Q1) and the highest single value), and the horizontal line at the end of the lower dashed whisker marks marks the lower extreme (the greater of Q1-1.5(Q3-Q1) and the lowest single value).

## DISCUSSION

Advances in high-throughput sequencing technology have made it possible to generate a wealth of viral genome information directly from clinical material. However, operational limitations should be registered in assessing the data. These include sample characteristics (source, viral content and presence of multiple strains), confounding factors (logistical errors and cross-contamination), design of the bait library and the sequencing protocol (ability to enrich all strains and acquire data across the genome), and quality and extent of the sequencing data (library diversity and coverage depth). Since perceived levels of intrahost variation are particularly sensitive to these factors, we proceeded cautiously with this aspect. However, as indicated in our earlier study [21], it is clear that the number of minor variants in single-strain infections was markedly less than that in multiple-strain infections. It was also far less than that reported by others in samples from congenital infections [16]. The factors listed above may have been responsible for the outliers observed in single-strain infections (Figure 2). In our view, accurate estimates of the levels of intrahost variation in single-strain infections are not available from the present and previous studies, and will require sequencing and bioinformatic approaches that are demonstrably reliable, robust and reproducible [46, 47].

Whole-genome analyses have confirmed the significant role of recombination during HCMV evolution reported in numerous earlier studies [12, 19]. Recombination has occurred over a very long period but nonetheless remains limited in extent, with surviving events being more numerous in long regions, less numerous in short regions, and rare or absent in hypervariable regions, as would be consistent with the role of homologous recombination. Recombination frequency is potentially restricted in some circumstances by functional interdependence within the same protein (e.g. gB) or possibly between separate proteins (e.g. gN and gO [25, 30, 43]). However, it is not known whether differential homologous recombination due to sequence relatedness is of general biological significance for the virus. Also, strains have circulated that seem not to have recombined over many thousands of years.

The extent to which recombinants arise and survive in individuals with multiple-strain infections is a separate topic. Except where populations fluctuate significantly and are sampled serially (e.g. RTR1 in collection 2), it is difficult to approach this using short-read data, as they are based on PCR methodologies prone to generating recombinational artefacts. Long- or single-read sequencing technologies and new bioinformatic tools are needed in this area. Also, conclusions drawn from transplant recipients, who are immunosuppressed and in whom HCMV populations may be diversified by transplantation from HCMV-positive donors or selected with antiviral drugs, are unlikely to represent naturally adapted cycles, such as those involving maternal transmission via breast milk [48].

The frequent identification of mutations giving rise to pseudogenes, and their apparently long history, reveals an interesting aspect of HCMV microevolution. The implication that some mutants have a selective advantage in certain circumstances may be extended to their evident ability to be transmitted congenitally and to be associated with neurological sequelae, probably in combination with specific host factors. The relevant genes are involved, or are suspected to be involved, in immune modulation. They include UL111A, which encodes viral interleukin-10 [49], UL40, which is involved in protecting infected cells against NK cell lysis [50] via its cleaved signal peptide, in which mutations occur, and UL9, which bears a potential immunoglobulin-binding domain [2].

Modern approaches offer a powerful means for analysing HCMV genomes directly from clinical material, with the important proviso that the data should be quality assessed and interpreted in the context of the known evolutionary and biological characteristics of the virus. Extensive high-throughput sequence data are likely to illuminate further the epidemiology, pathogenesis and evolution of HCMV in clinical and natural settings, thus facilitating the identification of virulence determinants and the development of new interventions.

## NOTES

### Acknowledgements

The authors thank Florent Lasalle, Daniel Depledge and Judith Breuer (University College London) for kindly providing unpublished collection 3 datasets and for updating the associated genome sequences in GenBank, and Jenny Witthuhn (Hannover Medical School) for excellent technical assistance.

### Supplementary data

Supplementary materials are available at The *Journal of Infectious Diseases* online. These consist of data provided by the authors to benefit the reader, are not copyedited, and are the sole responsibility of the authors. Questions or comments should be addressed to the corresponding author.

