## Supplementary Figure 1 for "Human Cytomegalovirus Genomes Sequenced Directly from Clinical Material: Variation, Multiple-Strain Infection, Recombination and Mutation"

**Supplementary Figure 1.** Nucleotide sequence alignments of genes used for genotyping or that commonly contain mutations.
Sequences are shown in the orientation present in the genome, with gaps introduced by alignment depicted by hyphens.
Sequence names are from Supplementary Table 1 or GenBank.
Genotype names are highlighted in green.

## A – RL5A

Initiation codons (the first and, where present, second ATG codons) and termination codons are highlighted in yellow. Sequences used for motif-matching are indicated in blue font (identifying genotypes) or red, gold or purple font (identifying G1 variants).
Cyan highlighting indicates strains in which this gene is mutated and the mutations responsible: substitutions that introduce in-frame stop codons (underlined) or insertions or deletions that cause frameshifts (the C-tract towards the right end of the alignment is not included).
The scattered locations of second ATG codons suggests that the true initiation codon for all genotypes is the first one. If this is the case, many more strains are mutated because of frameshifts in the C-tract, perhaps indicating that RL5A is essentially a nonfunctional gene that is in the process of being lost.

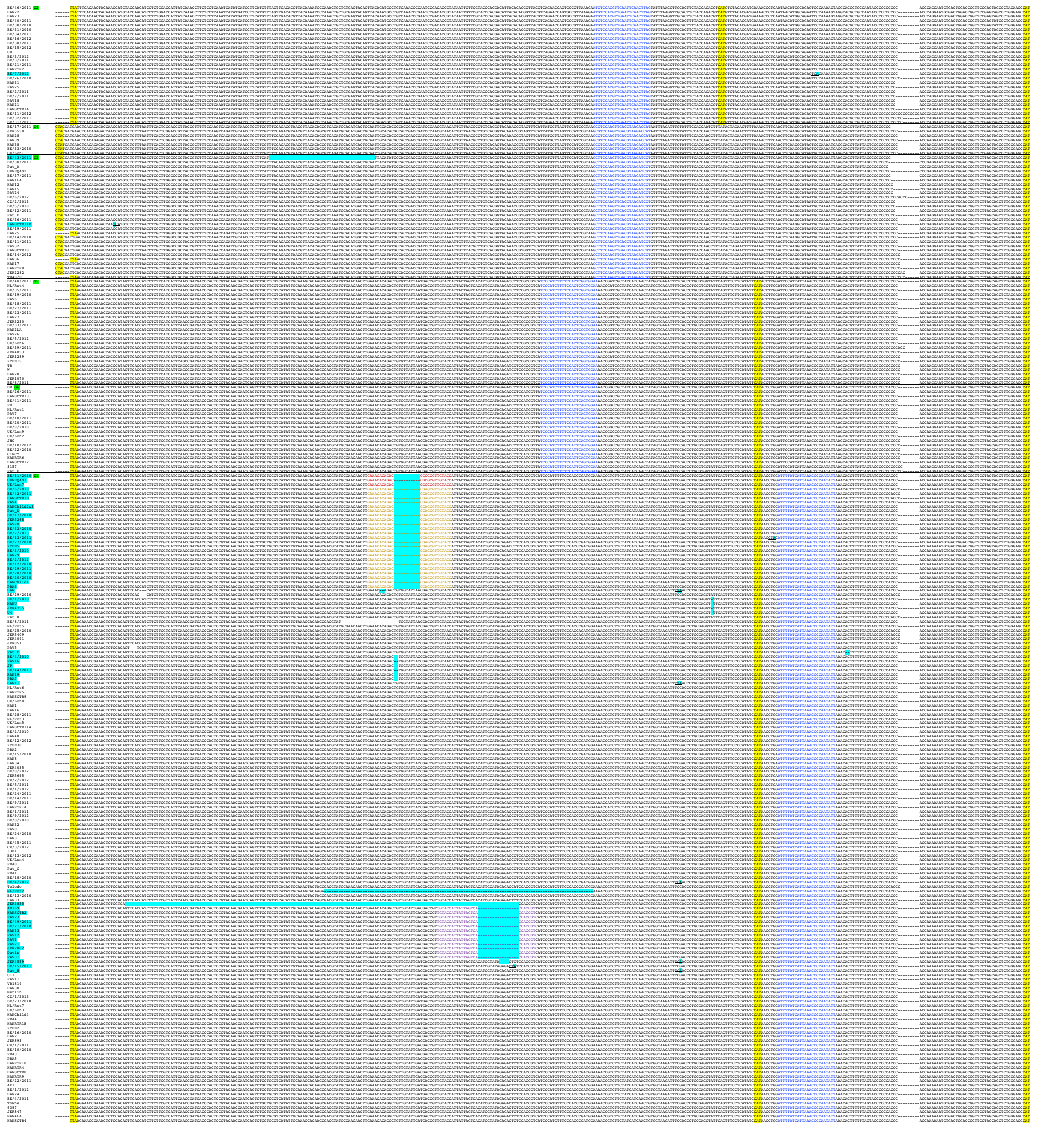

## B – RL6

Initiation codons (the first and, where present, second ATG codons) and termination codons are highlighted in yellow. Sequences used for motif-matching are indicated in blue font (identifying genotypes) or red font (identifying a common recombinant, with the region of recombination highlighted in grey).
Cyan highlighting indicates strains in which this gene is mutated and the mutations responsible: substitutions that introduce in-frame stop codons (underlined) or insertions or deletions that cause frameshifts (the C-tract towards the right end of the alignment is not included). The great majority of the gene is absent from G5.
The scattered locations of second ATG codons suggests that the true initiation codon for all genotypes is the first one. If this is the case, many more strains are mutated because of frameshifts in the C-tract, perhaps indicating that RL6 is essentially a nonfunctional gene that is in the process of being lost.

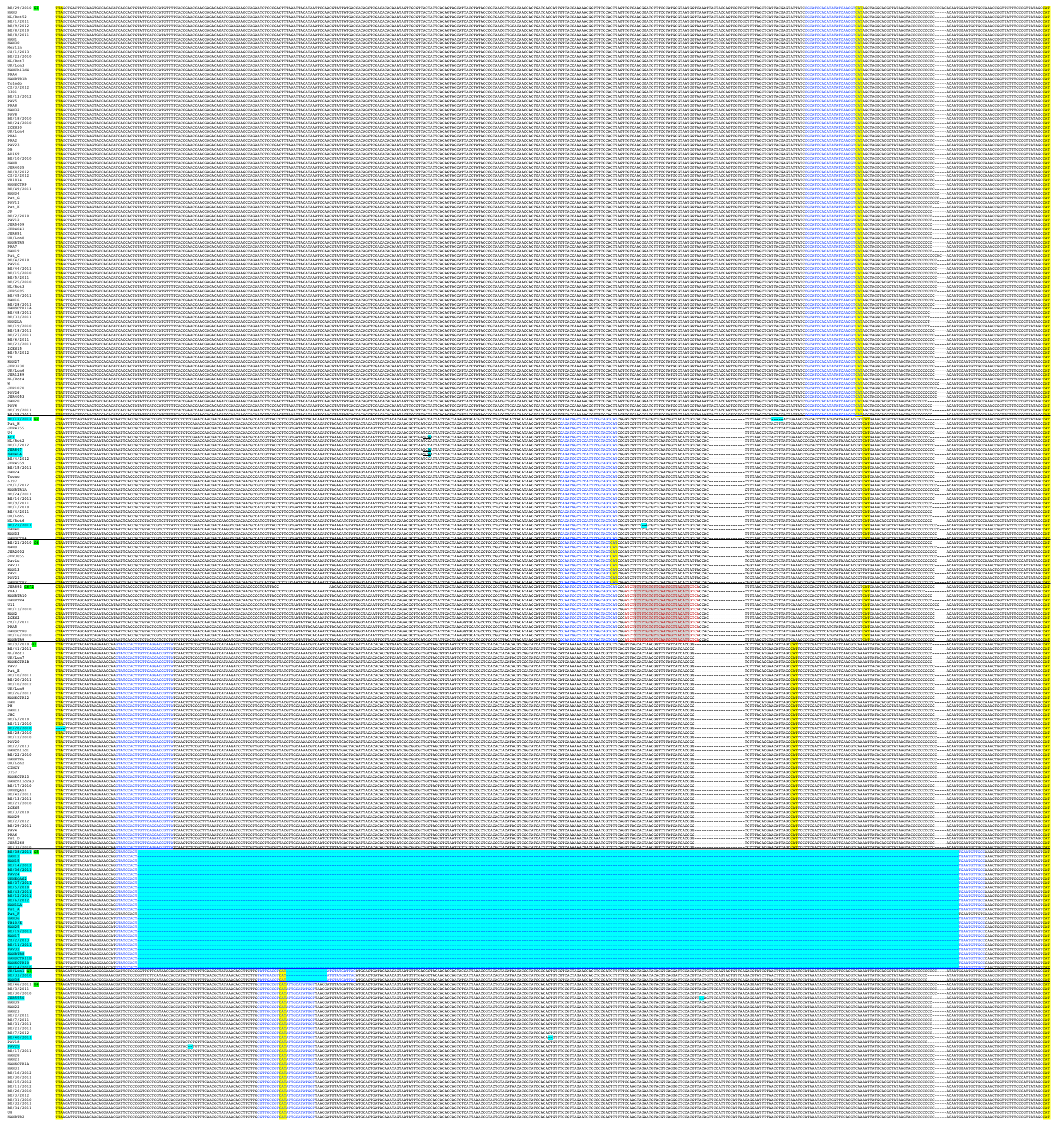

## C – RL12

Initiation and termination codons are highlighted in yellow. Sequences used for motif-matching are indicated in blue font (identifying genotypes) or red font (identifying a common recombinant, with the region of recombination highlighted in grey).
Cyan highlighting indicates strains in which this gene is mutated and the mutations responsible: substitutions that introduce in-frame stop codons (underlined) or insertions or deletions that cause frameshifts.

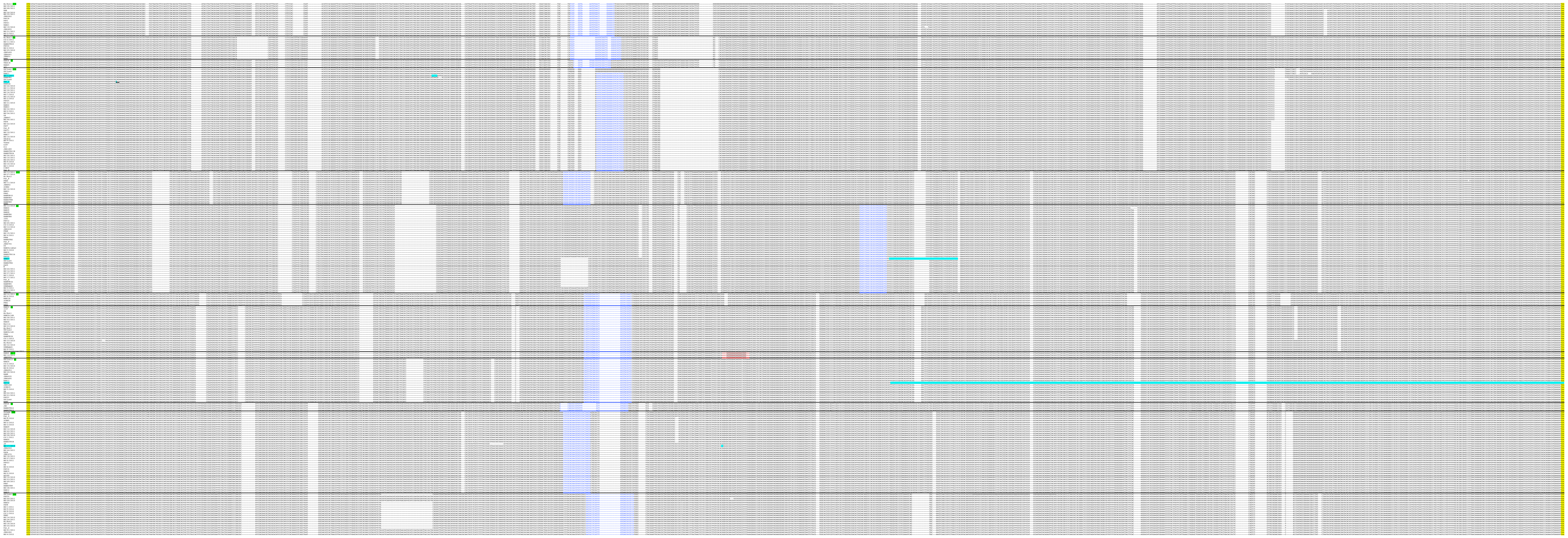

## D – RL13

Initiation and termination codons are highlighted in yellow. Sequences used for motif-matching are indicated in blue font (identifying genotypes).
Cyan highlighting indicates strains in which this gene is mutated and the mutations responsible: substitutions that introduce in-frame stop codons (underlined) or insertions or deletions that cause frameshifts. Mutations are likely not have been present in the clinical material but to have resulted during viral isolation in cell culture.

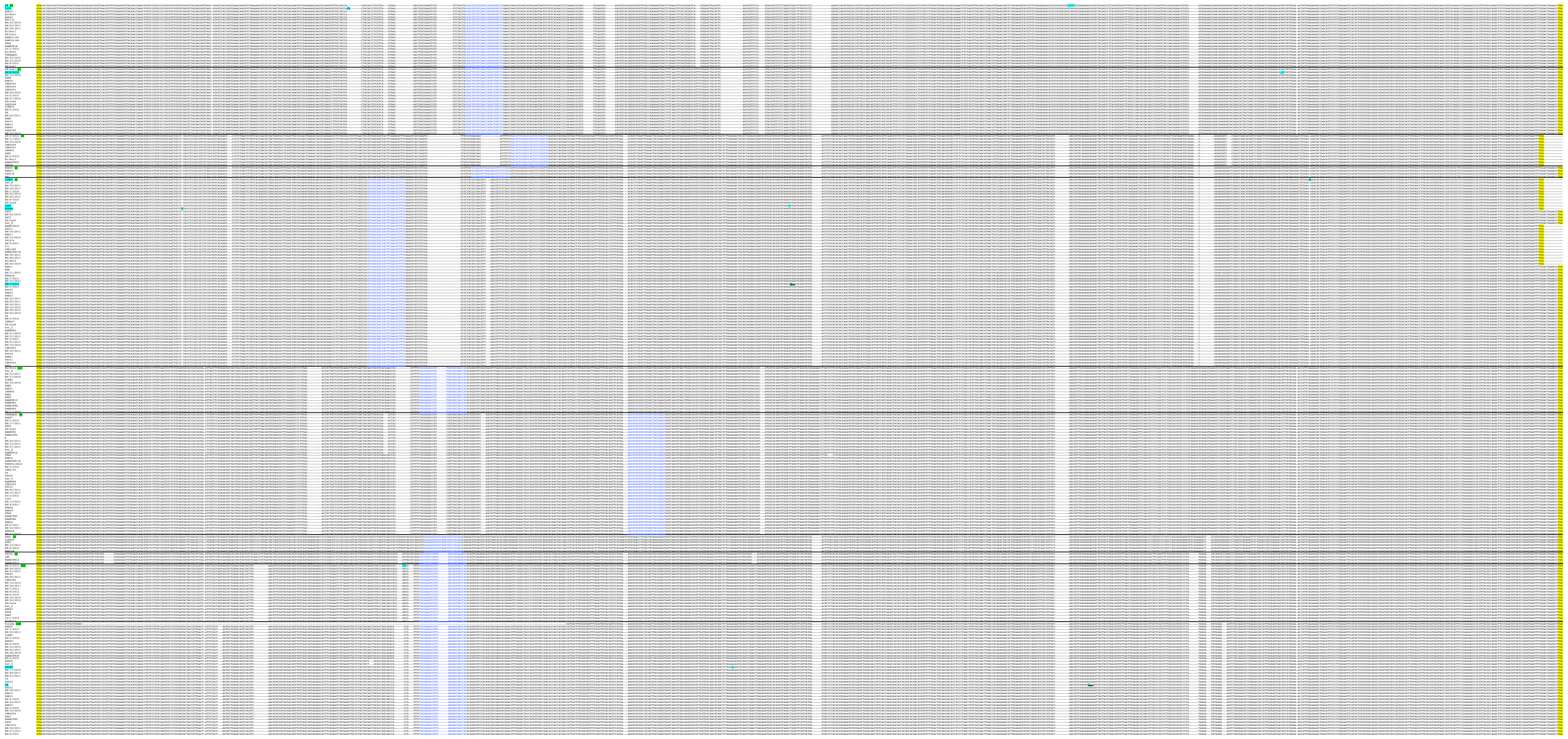

## E – UL1

Initiation and termination codons are highlighted in yellow. Sequences used for motif-matching are indicated in blue font (identifying genotypes).
Cyan highlighting indicates strains in which this gene is mutated and the mutations responsible: substitutions that introduce in-frame stop codons (underlined) or insertions or deletions that cause frameshifts. All G9 and G10 strains are mutants. It is not possible to discern the precise locations of the multiple mutations in G9.

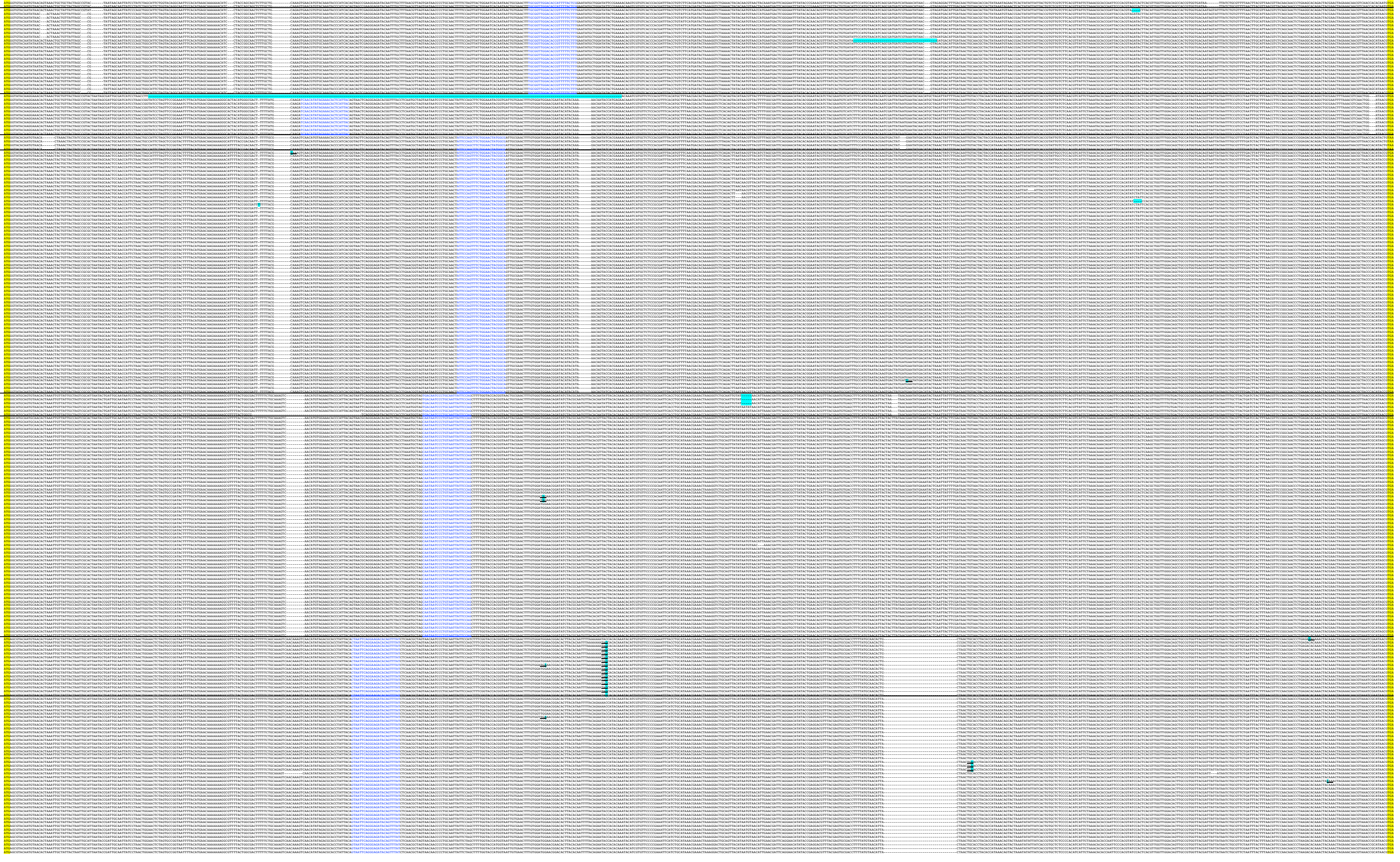

## F – UL9

Initiation and termination codons are highlighted in yellow. Sequences used for motif-matching are indicated in blue font (identifying genotypes).
Cyan highlighting indicates strains in which this gene is mutated and the mutations responsible: substitutions that introduce in-frame stop codons (underlined) or insertions or deletions that cause frameshifts. Mutations appear to be distributed more randomly in UL9 than in other genes, perhaps indicating that the selection pressures for loss of function are more recent.

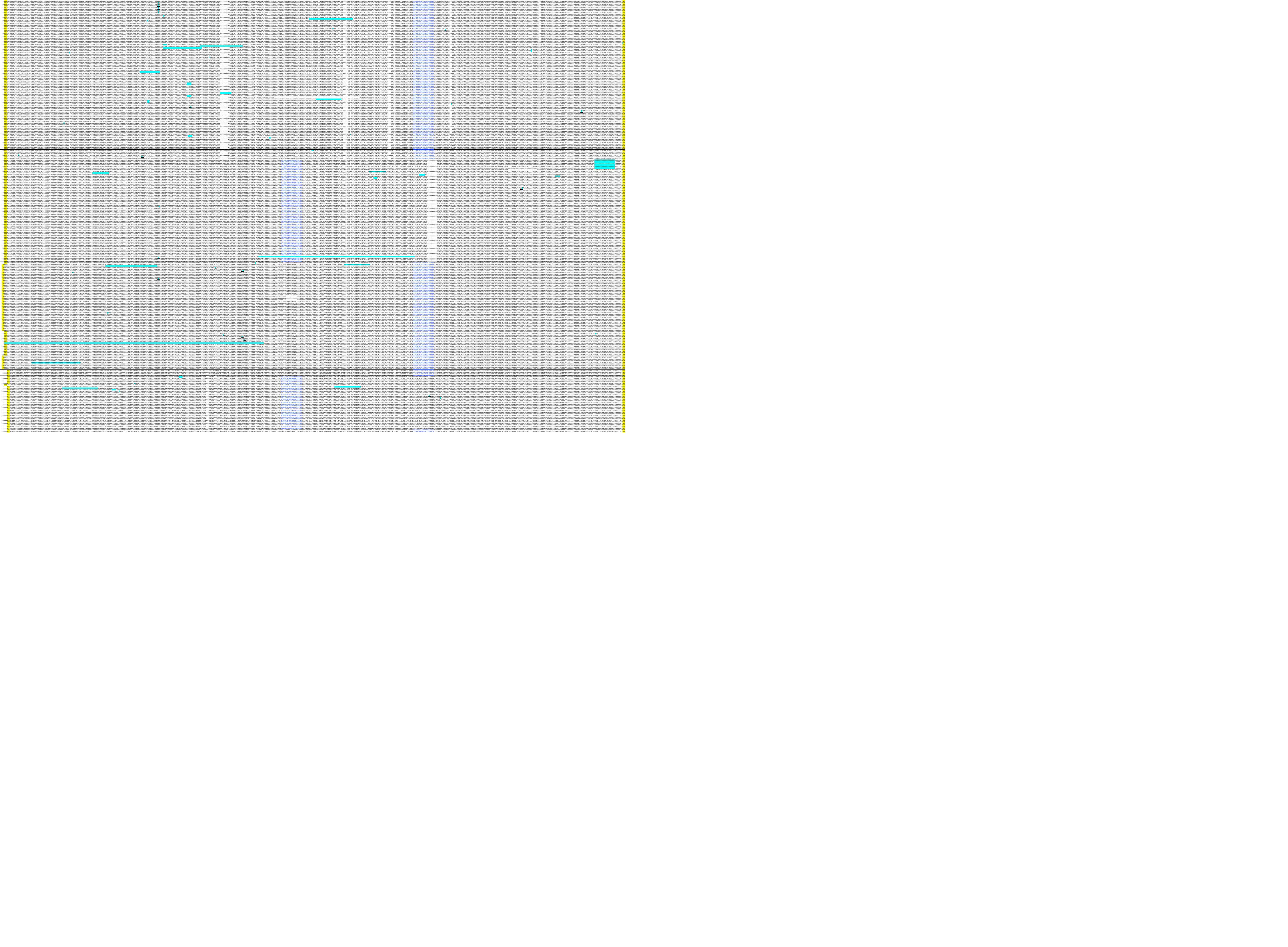

# G – UL11

Initiation and termination codons are highlighted in yellow. Sequences used for motif-matching are indicated in blue font (identifying genotypes).
Cyan highlighting indicates a strain in which this gene is mutated and the mutation responsible: a deletion that causes a frameshift.

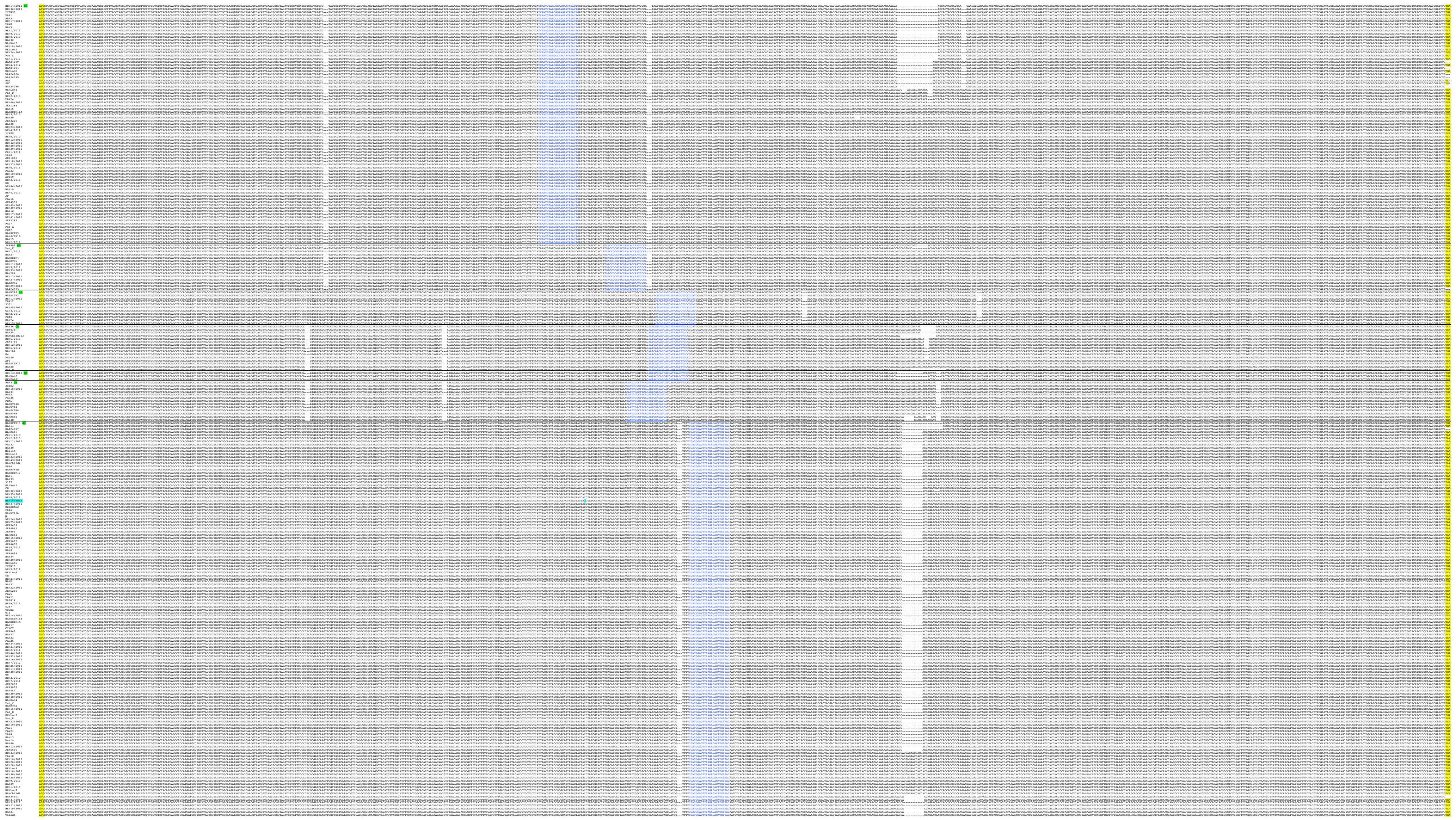

# H – UL20

Initiation and termination codons are highlighted in yellow.

### I – UL33 exon 2

Termination codons are highlighted in yellow. Exon ends are highlighted in magenta.

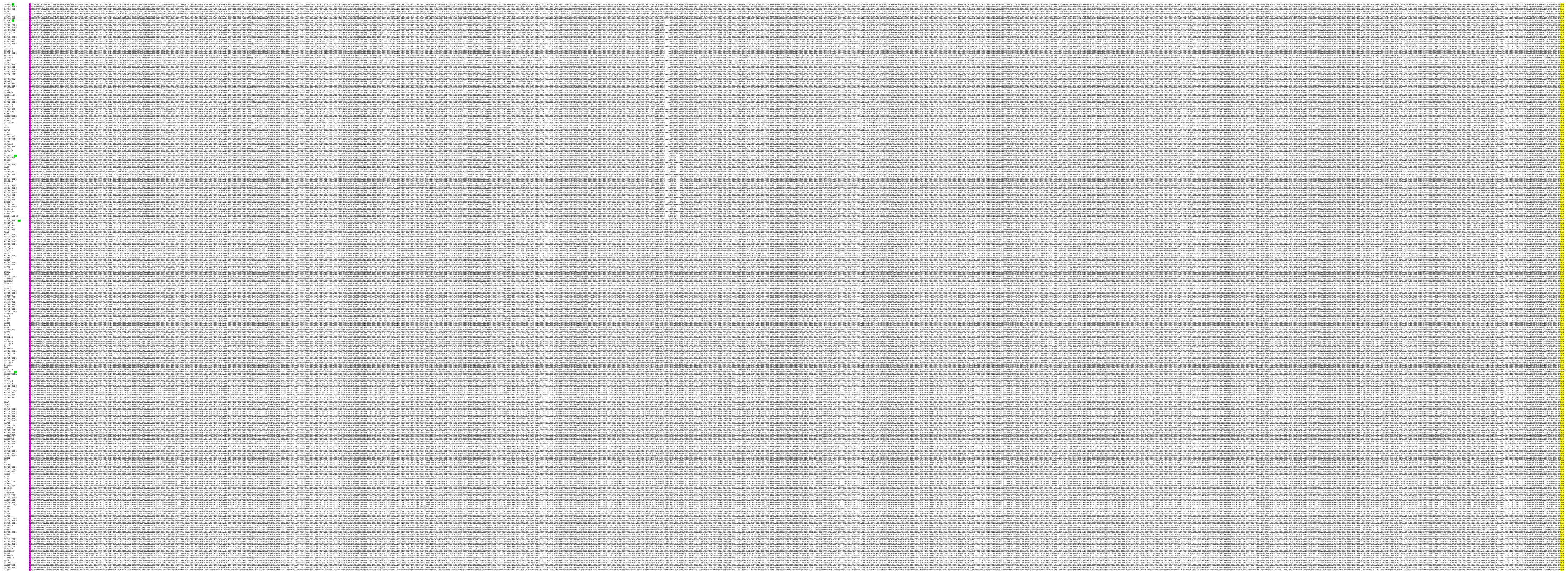

### J – UL37 exon 3

Termination codons are highlighted in yellow. Exon ends are highlighted in magenta.

[illegible]

## K – UL55N

Initiation codons are highlighted in yellow.

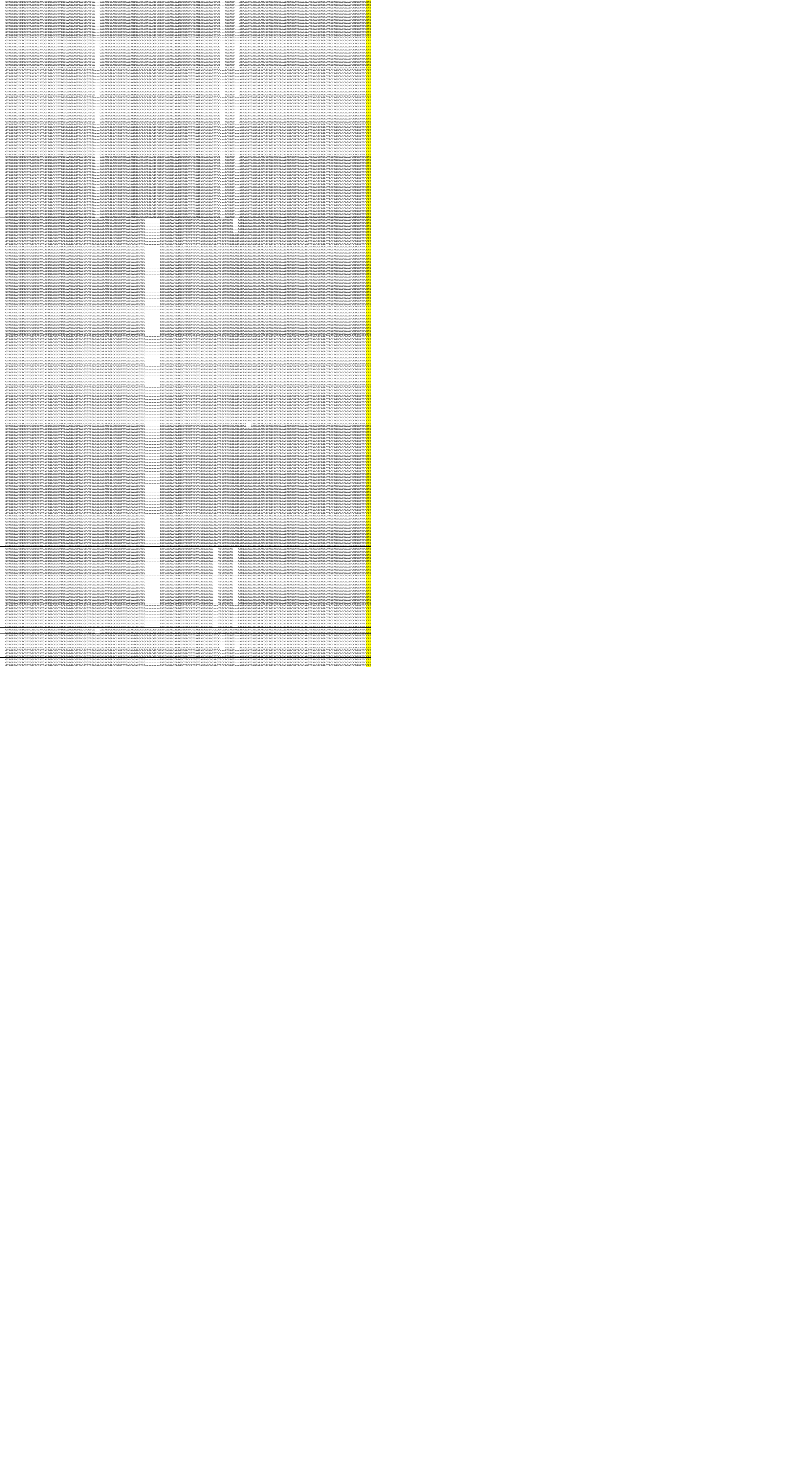

# L – UL73

Initiation and termination codons are highlighted in yellow. Sequences used for motif-matching are indicated in blue font (identifying genotypes). A recombinant is included.

[illegible]

# M – UL74

Initiation and termination codons are highlighted in yellow. Sequences used for motif-matching are indicated in blue font (identifying genotypes).

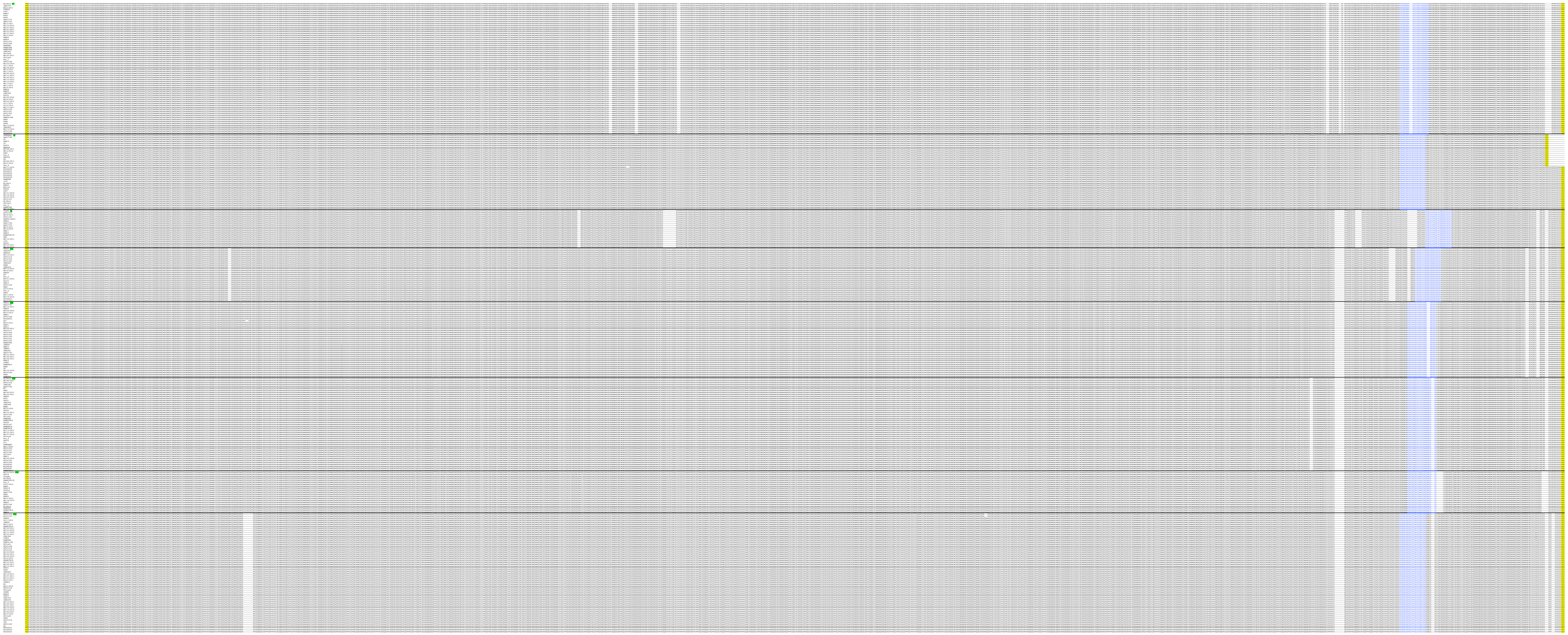

## N – UL120

Initiation and termination codons are highlighted in yellow. Sequences used for motif-matching are indicated in blue font (identifying genotypes).

[illegible]

# O – UL146

Initiation and termination codons are highlighted in yellow. Sequences used for motif-matching are indicated in blue font (identifying genotypes). Sequences corresponding to genotype-specific qPCR primers and probes are highlighted in green and grey, respectively.
The left primer had the sequence shown, and the right primer the complementary sequence. The probe had the complementary sequence, except for G3, G4, G5 and G14, in which it had the sequence shown.

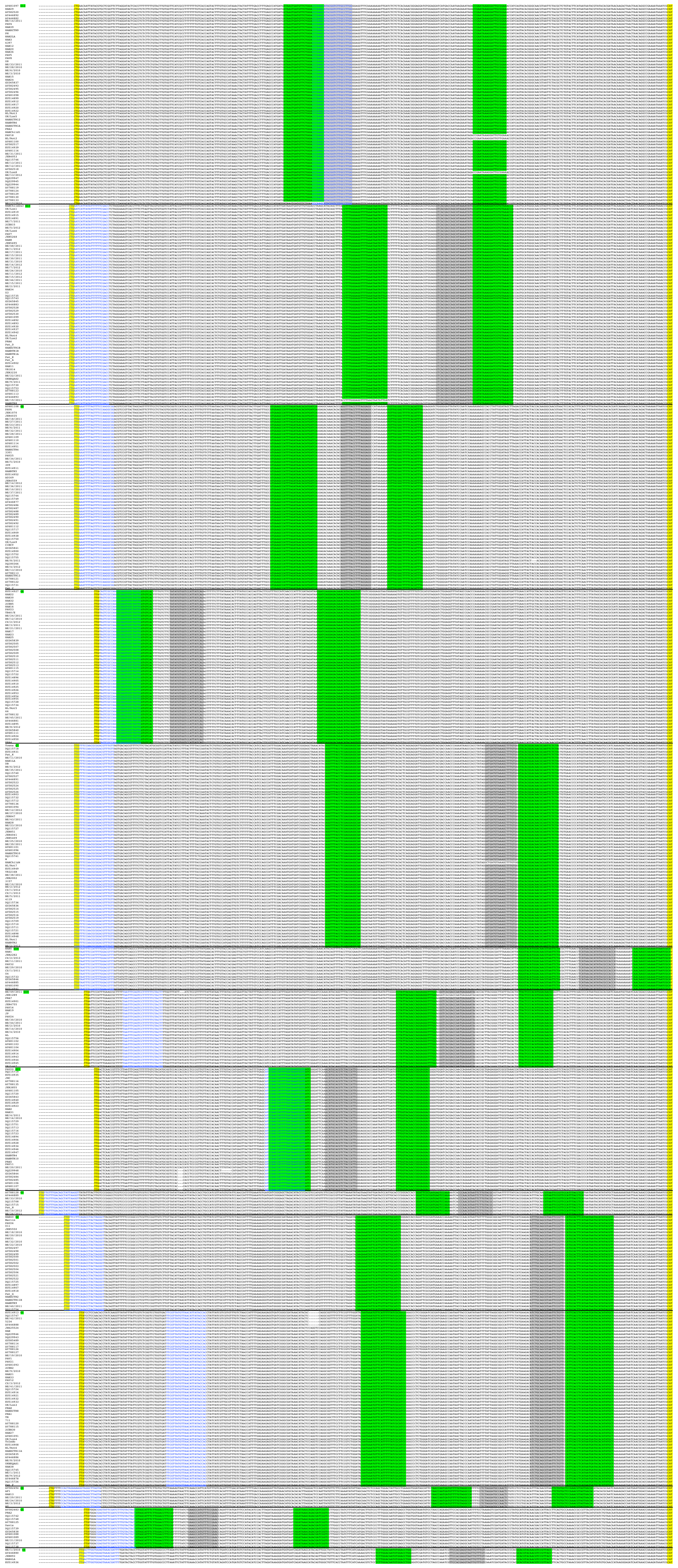

# P – UL139

Initiation and termination codons are highlighted in yellow. Sequences used for motif-matching are indicated in blue font (identifying genotypes). Two recombinants are included.

[illegible]

## Q – US9

Initiation and termination codons are highlighted in yellow. Sequences used for motif-matching are indicated in blue font (identifying genotypes) or red font (identifying a G2 variant).
Cyan highlighting indicates strains in which this gene is mutated and the mutations responsible: substitutions that introduce in-frame stop codons (underlined) or a deletion that causes a frameshift.

[illegible]

## R – UL111A

Initiation and termination codons are highlighted in yellow. Introns are highlighted in grey. This gene was not used for genotyping.
Cyan highlighting indicates strains in which this gene is mutated and the mutations responsible: substitutions that introduce in-frame stop codons (underlined) or insertions or deletions that cause frameshifts.

[illegible]
